# Serialized On-grid Lift-In Sectioning for Tomography (SOLIST)

**DOI:** 10.1101/2023.05.11.540146

**Authors:** Nguyen Ho Thuy Dung, Gaia Perone, Roberta Vazzana, Flaminia Kaluthantrige Don, Malan Silva, Simona Sorrentino, Paolo Swuec, Frederic Leroux, Nereo Kalebic, Francesca Coscia, Philipp S. Erdmann

**Affiliations:** Human Technopole, Milan, Italy; Leica Microsystems

## Abstract

Cryo-focused ion beam milling has enabled groundbreaking structural discoveries in native cells. Progress toward medically relevant applications, however, has been slow. We here present an adaptation of the cryo-lift out procedure for Serialized On-grid Lift-In Sectioning for Tomography (SOLIST), which increases throughput, reduces ice contamination, and enhances sample stability. With these improvements, new specimens, ranging from high-pressure frozen reconstituted LLPS droplets to human forebrain organoids, are accessible to cryo-electron tomography.

## Main Text

Compared to mechanical sectioning, gallium- and plasma-based focused ion beam (FIB) milling yield virtually artifact-free views of frozen-hydrated biological material. Lamellas, which contain cellular components of interest, can routinely be produced from plunge-frozen specimens and are thin enough for cryo-electron tomography (cryo-ET) and high-resolution subtomogram averaging (STA). While there is some concern regarding sample damage by FIB milling^1, 2^, *in situ* STA can reveal near atomic-level details of biological processes^3, 4^. To expand this concept of lamella milling to larger samples than single cells, high-pressure freezing (HPF) is required for vitrification^5^. HPF planchettes, however, are not compatible with standard on-grid milling procedures.

Cryo lift-out (LO) addresses this need by micromanipulators that operate with sub-micrometer precision to handle small portions of the high-pressure frozen material^6^. Adapted from the material sciences, cryo-LO grants access to biological material which may be too large for conventional lamella milling or the hybrid waffle method. Due to the more widespread availability and improved hardware, development has gained significant momentum over the past years^7, 8^. However, several aspects, including ease of operation, reproducibility, and sub-cellular precision targeting, still need to be improved so cryo-LO can become a routine method in structural biology.

Current best practices of the needle-based lift-out use a sacrificial adaptor made from copper (“adaptor chunk”) to attach the sample to the micromanipulator^9–11^. Although this improves success rates significantly, lift-in (LI), i.e., connecting the initial coarse lamella to the sample holder, requires specialized pin grids (Supporting Fig. 1a-b). While they are frequently used at room temperature, pin grids are problematic under cryo-conditions. First, lamellas are attached only to one side and break easily (Supporting Fig. 1c). Second, due to the lack of support, the drift of tomograms acquired far from the attachment site can be higher than those closer to the pin. While frame motion correction can compensate for sample movement, increasing stability and thereby minimizing drift already during data acquisition is preferable. To address this problem, custom-made slot-in grids, which support lamellas on both sides, have been explored in the past^7, 12^. However, their use is time-consuming and hard to realize with needle-based LO systems. Lastly, ice crystals frequently contaminate pin grid-mounted lamellas, which may be due to their exposed location and the higher electrostatic field strength on the pins (Supporting Fig. 1d). Based on these observations, ideal LI grids should be readily available, compatible with different chunk geometries and sizes, and guarantee sample integrity throughout the workflow.

For single particle analysis, grids made entirely from gold have proven helpful for high-resolution datasets^13^. The combination of mechanical stability, the matched expansion coefficient of grid and film, and electrical conductivity, which reduces beam-induced motion, make them ideal sample supports. Many of the same qualities would be desirable for lift-in grids.

To explore if holey gold films can also support LO lamellas, we lifted in chunks of 25 µm x 10 µm x 3 µm and larger to all-gold grids under shallow angles and fixed the pieces on the film using micro-sutures (Supporting Fig. 2a-e). After clearing small areas in front of the coarse lamellas to prevent uncontrolled ripping of the gold film (Supporting Fig. 2b), we polished the leading edges and coated them with the gas injection system (GIS) to reduce curtaining (Supporting Fig. 2c). In the final step samples were milled analogously to cells on grids, yielding lamellas transparent to transmission electron microscopy (TEM) (Supporting Fig. 2f-i), confirming that commercially available gold grids are stable enough to support LI applications.

Since the rate-limiting step in cryo-LO is preparing the lift-out site, it is wasteful only to cut a single lamella from the material. To overcome this limitation, we devised a method to enhance throughput by dividing the initial sample chunk into sections of 2-4 µm each, which we placed on individual grid squares (Fig. 1a). Each time, a cut of 1-2 µm was made to release the lamella from the remaining lift-out material so that in total each section requires about 4-5 µm of the initial chunk. Using this sequential LI, a 25 µm-tall piece is split into four to six individual lamellas (Fig. 1b-c, Supporting Video 1). This Serialized On-grid Lift-In Sectioning for Tomography (SOLIST) significantly improves sample use, as several lamellas are obtained from the same cell, small organism, organoid, or tissue biopsy. SOLIST effectively multiplies the number of LO lamellas produced in each session (Supporting Table 1, Supporting Video 2). A similar concept that relies on lifting in chunks on bare grids without film support was published in parallel to our procedure^14^.

**Figure 1.**
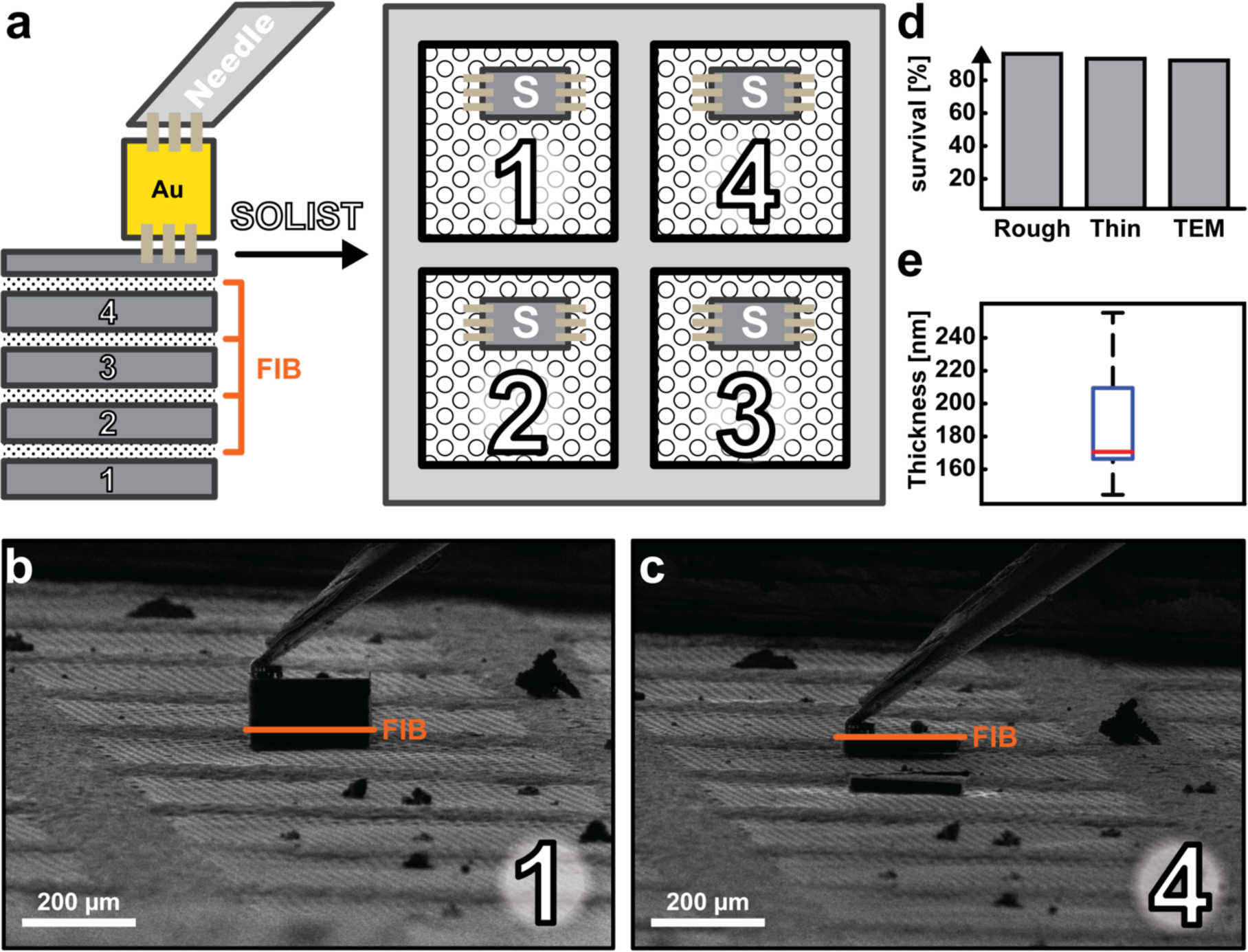
Serialized On-grid Lift-In Sectioning for Tomography (SOLIST). **a,** Schematic representation of the SOLIST procedure. **b-c,** FIB view images of individual SOLIST slices attached to the grid. **d,** Success rates of SOLIST lamellas milling by step. **e,** Box plot of SOLIST lamella thickness. Mean ± SD: 187 ± 31 nm (n =; whiskers indicate maximum and minimum values, the box top the 75^th^ percentile and the bottom the 25^th^ percentile; red line indicates the median).

Lamellas prepared with SOLIST are highly stable during storage, transfer to the TEM, and can be subjected to several loading/unloading cycles, for instance, for cryo-CLEM applications (Fig. 1d). Their thickness is comparable to regular on-grid lamellas (Fig. 1e) and contamination with ice crystals is considerably reduced compared to lift-out on pin grids (Supporting Fig. 2f-i).

To probe the generalizability of our method, we tested two representative yet vastly different scenarios: 1) *in vitro* samples of reconstituted liquid-liquid phase separated (LLPS) chromatin droplets and 2) forebrain organoids, which are particularly interesting as medically relevant model systems to study various neurodevelopmental and neuropsychiatric diseases^15–19^.

In general, SOLIST can be applied to any sample amenable to HPF and does not distinguish between planchettes and waffles for the LO chunk preparation (Fig. 2a, Supporting Fig. 3). While a part of the lift-out material is required for attachment and support on the grid (∼ 4-5 µm on either side), sufficiently large lamellas (15-25 µm x 20 µm) can regularly be produced. Furthermore, since they are highly stable and their geometry is well-defined, it is straightforward to automate parts or the entire milling process using available software packages^20, 21^.

**Figure 2.**
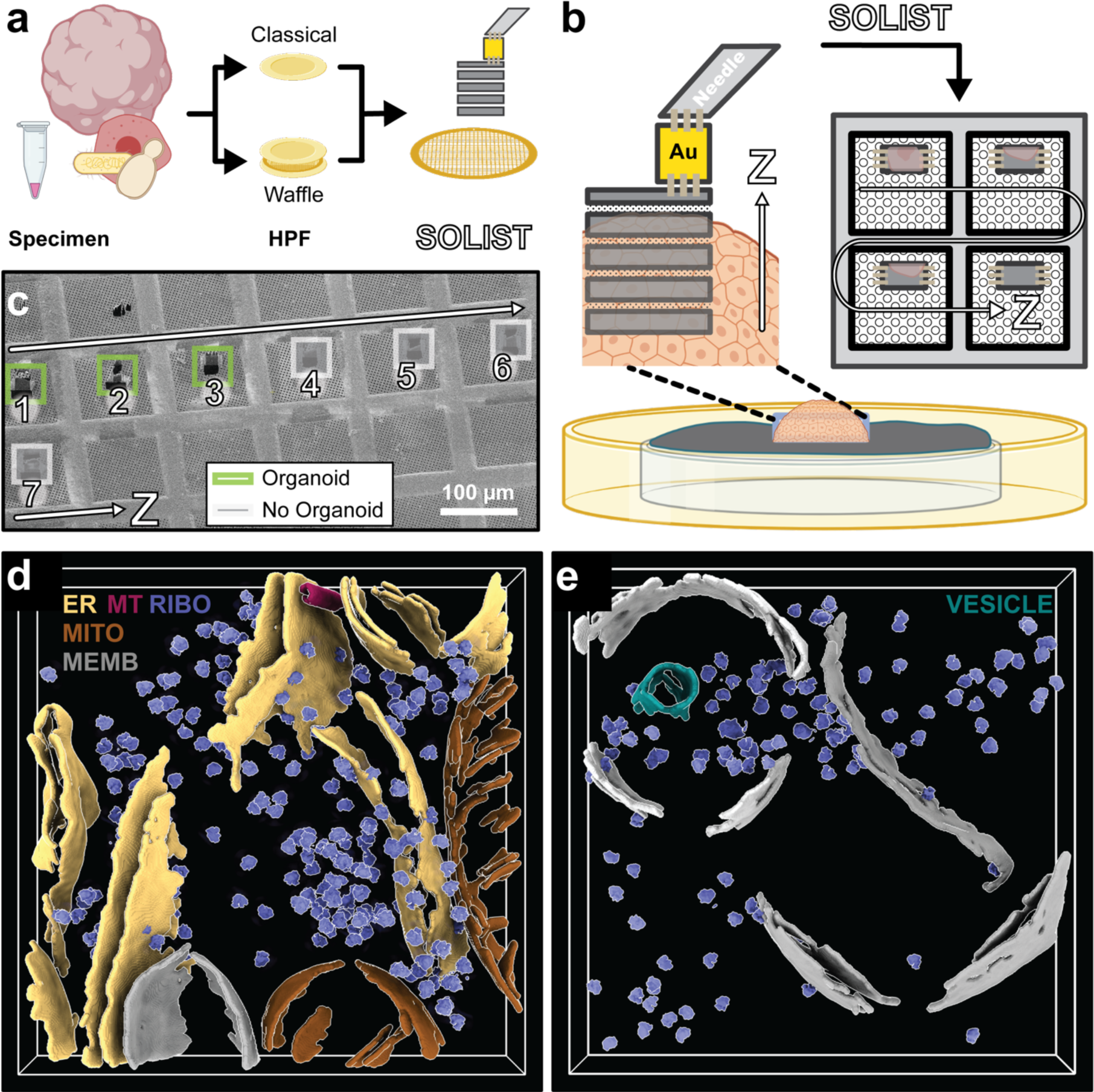
SOLIST enables investigations in complex samples, including developing forebrain organoids. **a,** Generalized workflow from sample to SOLIST. **b,** A chunk of HPF material is translated into a series of LO slices on a gold grid. The lack of accuracy in z-targeting is compensated by the horizontal series. **c,** Within the series of lamellas, organoid material is found in the deeper (i.e. earlier) layers. **d-e,** Segmentations of representative brain organoid tomograms from adjacent SOLIST sections. ER: endoplasmic reticulum; MITO: mitochondria; MEMB: membrane; MT: microtubule; RIBO: ribosome. (See Supporting Video 3).

For high-content samples (e.g. cell suspensions), LO sites can be prepared without targeting. However, an additional correlation step is required to identify the target region for organoids. To this end, reflected light and fluorescence overviews are acquired directly on the high-pressure frozen sample. With this information, areas of interest are localized in x and y, before LO sites are prepared using the focused ion beam. As an alternative to site preparation by milling, planchettes may be trimmed (planed) on a cryo-ultramicrotome to reduce the overall FIB time (Supporting Fig. 4).

Using the sequential lift-out approach, deeply submerged biological structures can be excavated from the planchette, compensating for the lack of targeting in z (Fig. 2b-c). In general, SOLIST yields relatively large lamellas. Hence, high-throughput approaches for cryo-ET data acquisition may be used to optimize microscope use^22, 23^. Tomograms of our lamellas produced high-resolution snapshots of the interior of forebrain organoids and their molecular contents (Fig. 2d-e, Supporting Videos 3-4), while previous work was limited to imaging only the periphery, thin enough for direct investigation by transmission electron microscopy ^24^.

Considering the stability of coarse lamellas on gold grids, we reasoned that targeting in HPF samples could further be refined by 3D correlation. SOLIST reduces several problems commonly associated with correlative focused ion beam milling: 1) lift-in on clean grids reduces ice contamination which can interfere with detecting fiducial beads; 2) the well-defined geometry of the lamellas, and especially their initial thinning to 3-4 µm, reduces fluorescence imaging artifacts that arise from the refractive index mismatch between the chunk and its surrounding; 3) since HPF samples can be prepared with higher densities of target structures, the chance of finding the area of interest in the final lamella is significantly increased.

Based on this idea, we devised a two-step procedure for 3D correlation on HPF samples containing LLPS droplets. First, initial lamellas are prepared as described above and placed on gold grids coated with fiducial beads for correlation (Fig. 3a).

**Figure 3.**
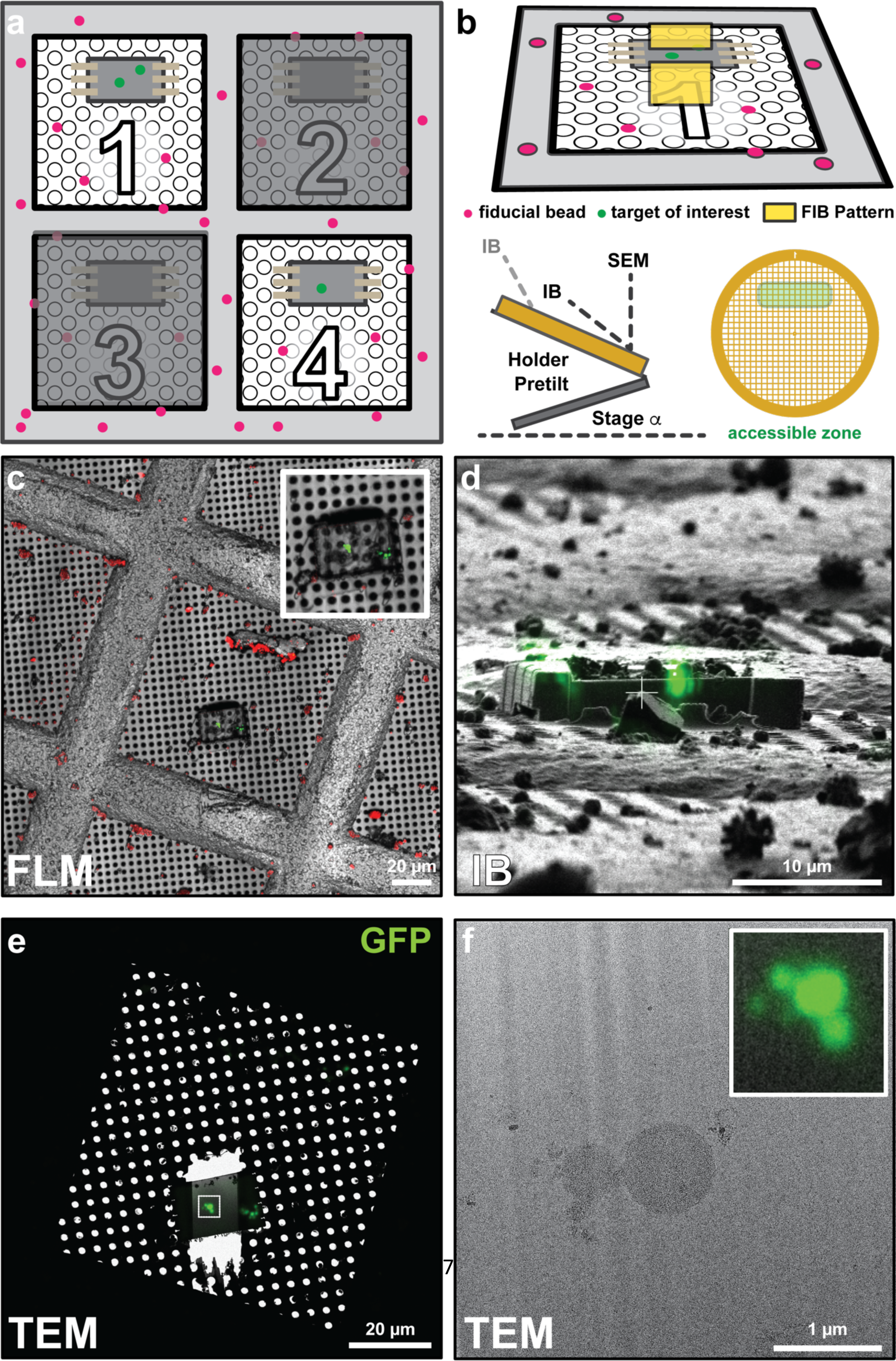
SOLIST allows 3D-correlative targeting in HPF samples. **a,** Initial SOLIST lamellas are screened for the target of interest (square 1 and 4). **b,** Concept of 3D correlation on a coarse lamella showing fiducial beads and overlay with the fluorescence data rotated to match the view of the ion beam (IB). Due to the shallow milling angle, only an array of 4×12 squares on the gold grid is accessible for lift-in and correlation. **c,** Scanning electron microscope (SEM) view overlaid with FLM data (green: LLPS droplets, red: fiducial beads) before milling. **d,** 3D CLEM on the coarse SOLIST chunk using the registration of the fiducial beads to transform the FLM data. **e,** TEM overview overlaid with a fluorescence image. **f,** Zoomed-in TEM view of the lamella in Fig. 3e (Insert: FLM data from Fig. 3e).

After transfer to the fluorescence microscope, widefield or confocal z-stacks are acquired depending on the required targeting accuracy before loading the sample back on the FIB, where 3D correlative milling is performed analogously to cells on grids (Fig. 3b)^25, 26^. The defined geometry of the coarse lamellas is particularly well suited for FIB automation after the lamella positions have been transferred. This ensures the best performance of 3D-targeted milling, and that the region of interest is contained in the final lamella (Fig. 3c-f).

In summary, SOLIST allows efficient preparation of several cryo-FIB lamellas from the same high-pressure frozen sample and with fluorescence-guided targeting. This includes, for the first time, detailed inner views of developing brain organoids. Our new approach improves targetability, throughput and lamella stability compared to previous implementations. While SOLIST is still not as efficient as traditional serial section approaches at room temperature, where layers of 70-100 nm can be cut, it represents a significant improvement over previous cryo-LO protocols. With critical bottlenecks of cryo-lift-out taken care of, we anticipate cryo-electron tomography datasets soon being generated for many new sample types such tissues and patient biopsies. Access to this high-resolution information will no doubt help advance our molecular understanding of healthy and pathological molecular processes and be a first step towards a true “biopsy at the nanoscale”.

## Acknowledgements

We thank Alessandro Vannini, Gaia Pigino, and Ana Casañal for helpful insights and discussion; Nikolai Klena, Helen Foster, and Sam Lacey for valuable comments; Daniel Panne and Ziad Ibrahim for providing the *in vitro* chromatin samples and the Cryo-Electron Microscopy Facility of Human Technopole for technical support and assistance. This work was supported by core funding from the Human Technopole. The Coscia lab is supported by the European Research Council (ERC-2021-STG Thyromol #101041298).

## Author contributions

NHTD, GP: Conceptualization, Methodology, Validation, Investigation, Data Curation, Resources, Writing - Review & Editing, Visualization; RV: Investigation, Resources, Writing - Review & Editing, Visualization; FKD: Investigation, Resources; MS: Investigation, Resources; SS: Investigation, Resources; PS: Investigation, Resources, Writing - Review & Editing; FL: Investigation, Resources; NK: Resources, Supervision, Funding acquisition, Writing - Review & Editing; FC: Resources, Supervision, Funding acquisition; PSE: Conceptualization, Methodology, Software, Validation, Investigation, Data Curation, Resources, Writing - Original Draft, Writing - Review & Editing, Visualization, Supervision, Funding acquisition

## SUPPORTING MATERIAL

**Supporting Table 1.** Typical milling time for the SOLIST approach per step.

|  | *Step* | *Current* | *Timing* |  |
| --- | --- | --- | --- | --- |
| LO Type |  |  | *From milling direction* | *From perpendicular direction* |
| *Sample trench* | Site preparation | 5 nA | 4 min<br>+ 10 min cleaning | 7 min<br>+ 10 min cleaning |
|  | Surface cleaning | 300 pA | 5 min | 10 min |
|  | Leading edge polishing | 300 pA | 5 min |  |
|  | Undercut milling | 500 pA | 10 min | 15 min |
|  | Sample Detachment | 300 pA | 3 min |  |
|  |  |  | **TOTAL / Site** |  |
|  |  |  | 37 min | 50 min |
| *Sample section* | SOLIST Section Attachment | 1 nA | 3 min / Section |  |
|  | Attaching from top | 30 pA | 1 min / Section |  |
|  | Leading edge polishing | 300 pA | 2 min / Section |  |
|  | Surface cleaning | 300 pA | 5 min / Section |  |
|  |  |  | **TOTAL / Section** |  |
|  |  |  | 11 min |  |

**Supporting Figure 1.**
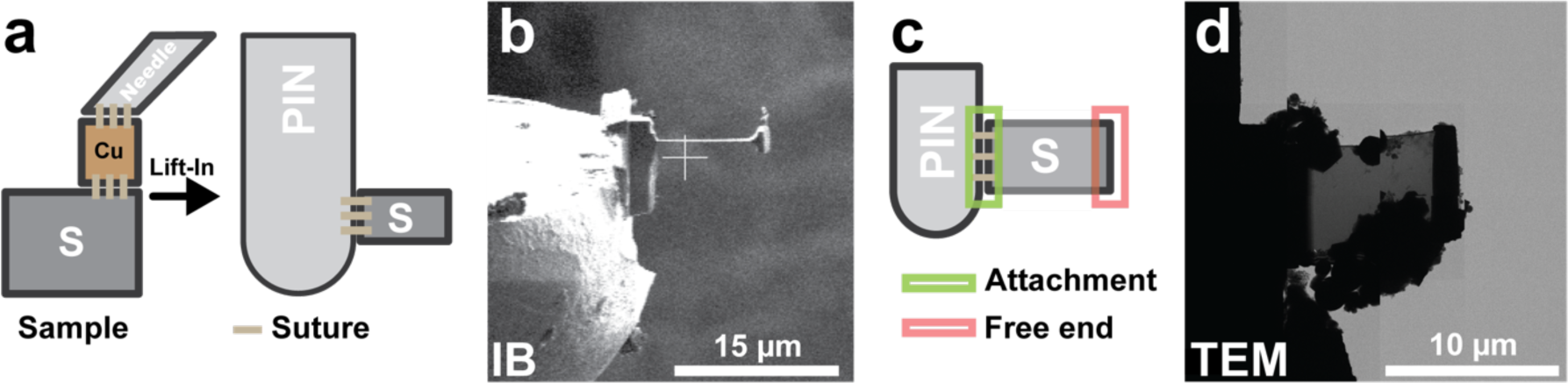
Traditional lift-in procedure on pin grids. **a,** Schematic of the conventional lift-in procedure. **b,** FIB view of a single LO lamella attached to a pin grid. **c,** Conventional sample attachment results in lamellas supported only on one side. **d,** Example of ice crystals-contaminated lamella on a pin grid (TEM montage).

**Supporting Figure 2.**
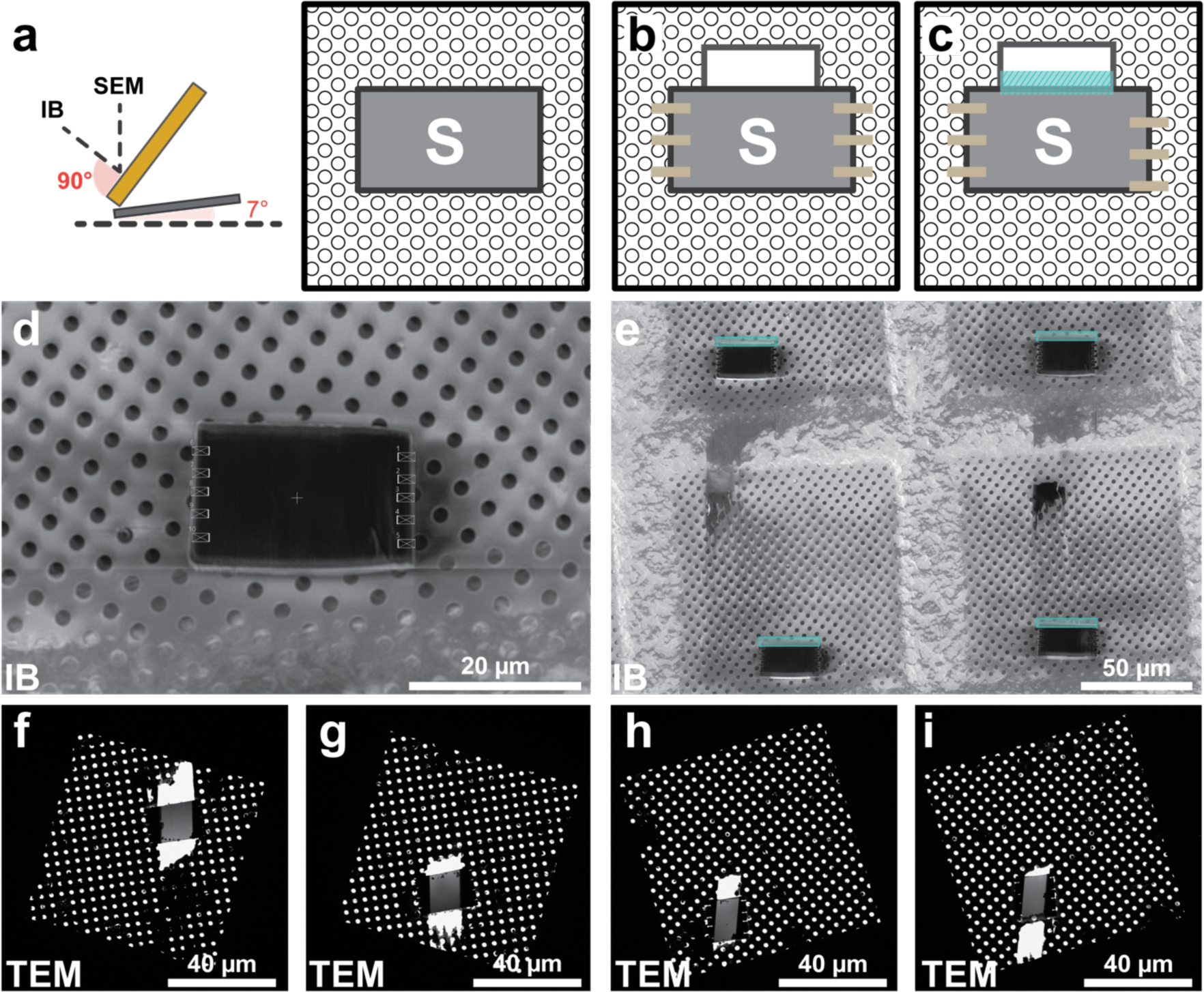
Lift-in on gold grids using SOLIST procedure. **a,** With the coarse sections attached on the EM grid, the stage is rotated to the perpendicular position (stage tilt = 7°, stage rotation = 0°). **b,** From this position, parts of the gold film are removed to prevent ripping and micro-stitches are milled with cleaning cross-section patterns at 30 pA. The milling direction is towards the center of the section, and z dimension is set to 0.5 μm. **c,** The leading edge is cleaned with a cleaning cross-section at 300 pA. **d,** FIB image of a coarse SOLIST section with the patterns used for attachment. **e,** A set of SOLIST sections after attachment. Light-blue rectangles indicate the leading edges. **f-i,** TEM montages of electron-transparent lamellas show low ice contamination.

**Supporting Figure 3.**
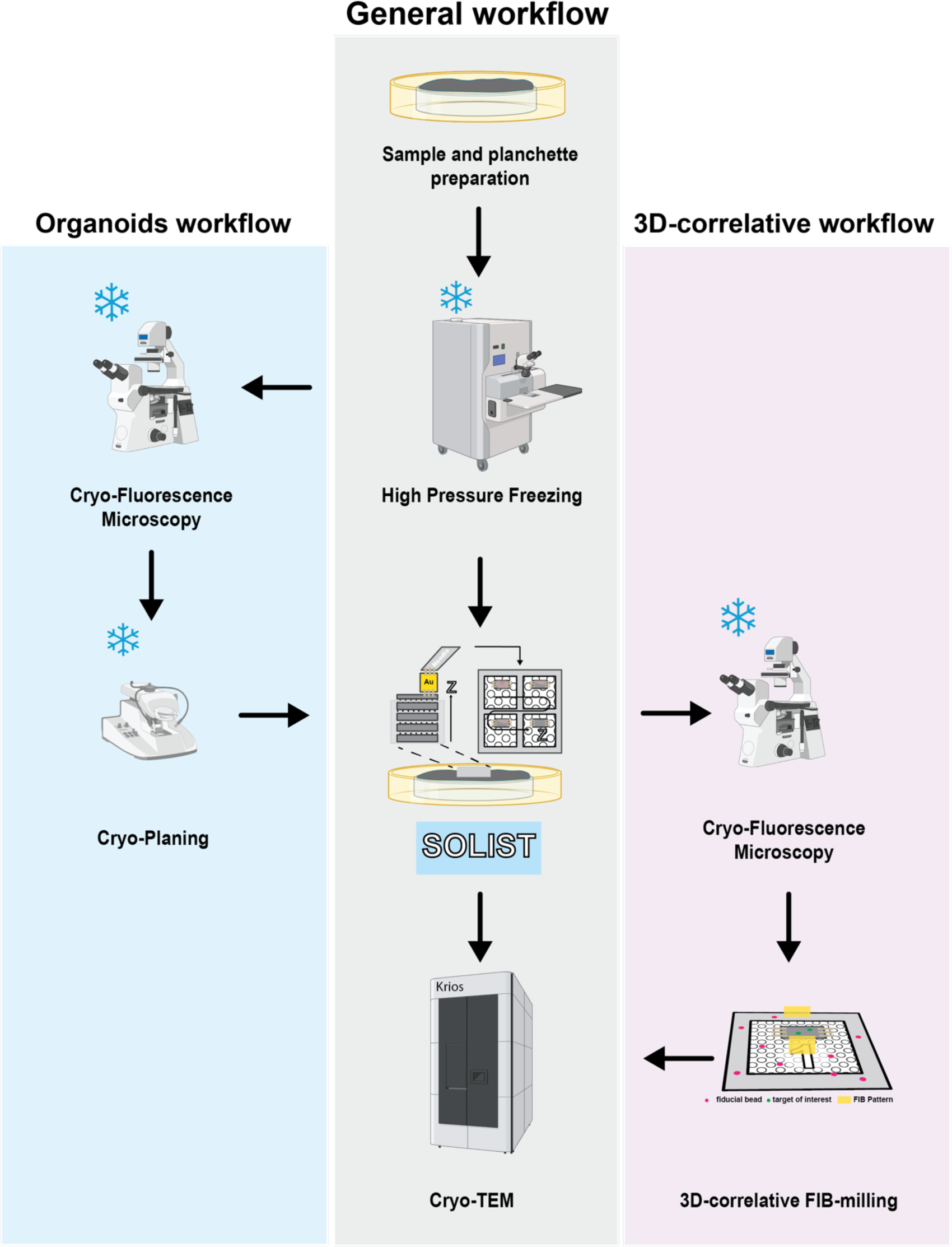
SOLIST for *in situ* cryo-ET of HPF-frozen samples. General workflow for the SOLIST procedure. For organoid samples, imaging and buffer removal by cryo-planing can be performed before SOLIST to expose the organoid to the surface. For the 3D-correlative workflow, target positions in coarse sections are identified by fluorescence microscopy before fine milling.

**Supporting Figure 4.**
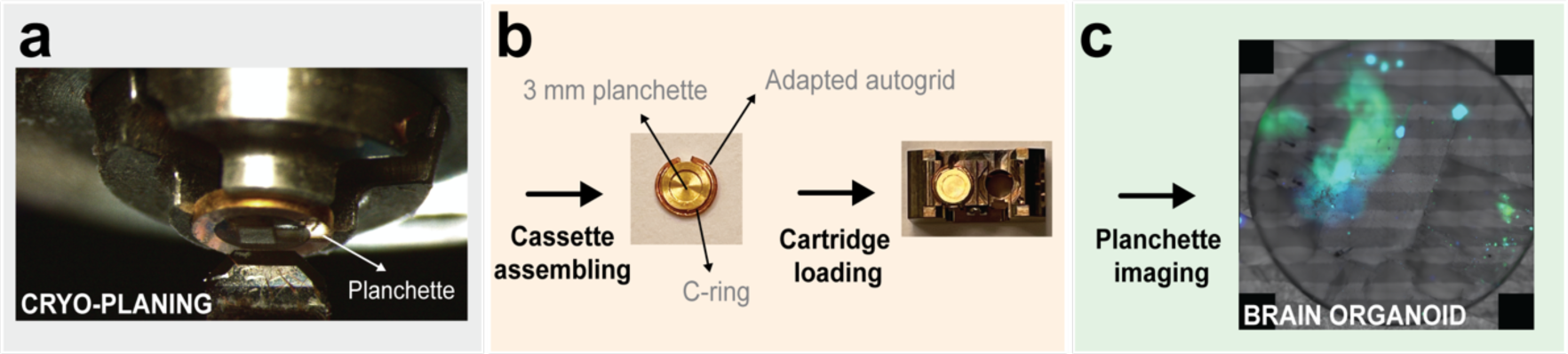
Detailed steps for the organoid workflow. **a,** Cryo-planing of HPF planchettes is performed to expose the organoid. **b**, A customized adaptor, composed of a cut autogrid and a C-ring, is used to load the planchette in the cryo-fluorescence microscope cartridge. **c**, Representative overview of a HPF planchette containing brain organoids. Fluorescent channels overlayed with reflected light (Green: GFP; Blue: Hoechst).

**Supporting Figure 5.**
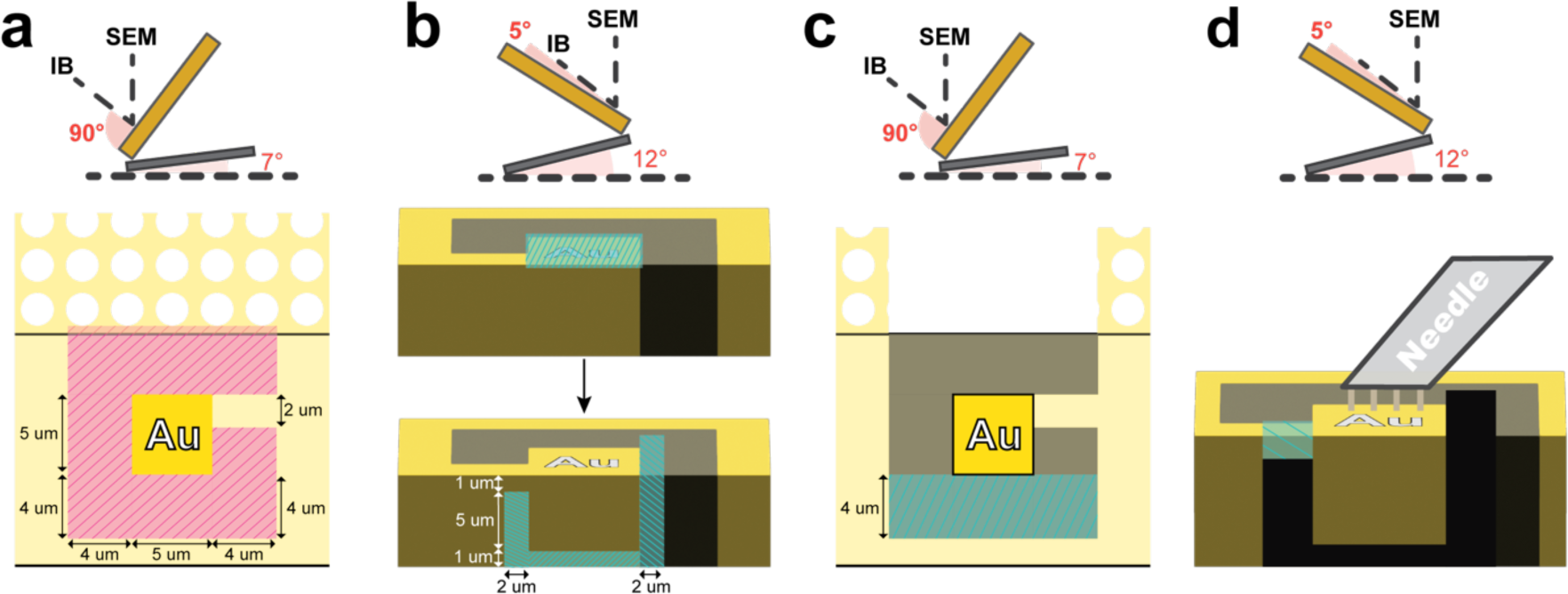
Gold adaptor chunk preparation. **a**, Site preparation from the perpendicular position. **b**, Surface cleaning and undercut milling from the milling position at 12° stage tilt. **c**, Redeposition cleaning from the perpendicular position. **d**, Needle attachment from the milling position at 12° stage tilt.

**Supporting Figure 6.**
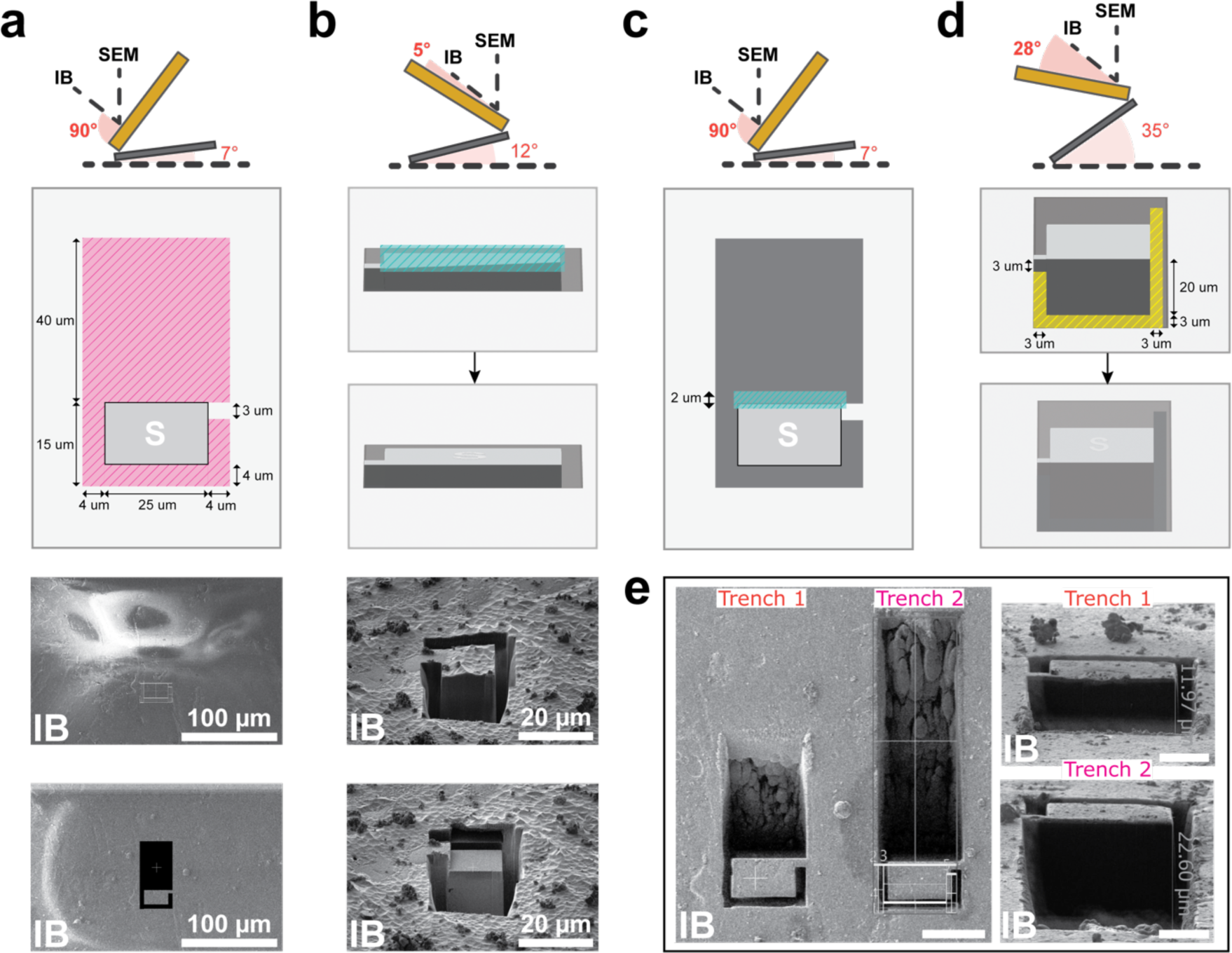
Sample site preparation for LO from the milling direction. **a**, Sample trench preparation from the perpendicular position. At the bottom, FIB images of the frozen sample with the patterns used to prepare the chunk site from the perpendicular position before and after milling. **b**, Surface cleaning for needle attachment from the milling position at 12° stage tilt. At the bottom, FIB images of the frozen sample before and after milling. **c**, Leading edge polishing from the perpendicular position. **d**, Undercut and sides cleaning from the milling position at 35° stage tilt. **e**, The amount of material cleaned in the front of the LO site defines the dimension of the sample chunk accessible for LO. The undercut angle is linked to the length of the cleaning trench. Scale bars represent 25 µm (left panel) and 10 µm (right).

**Supporting Figure 7.**
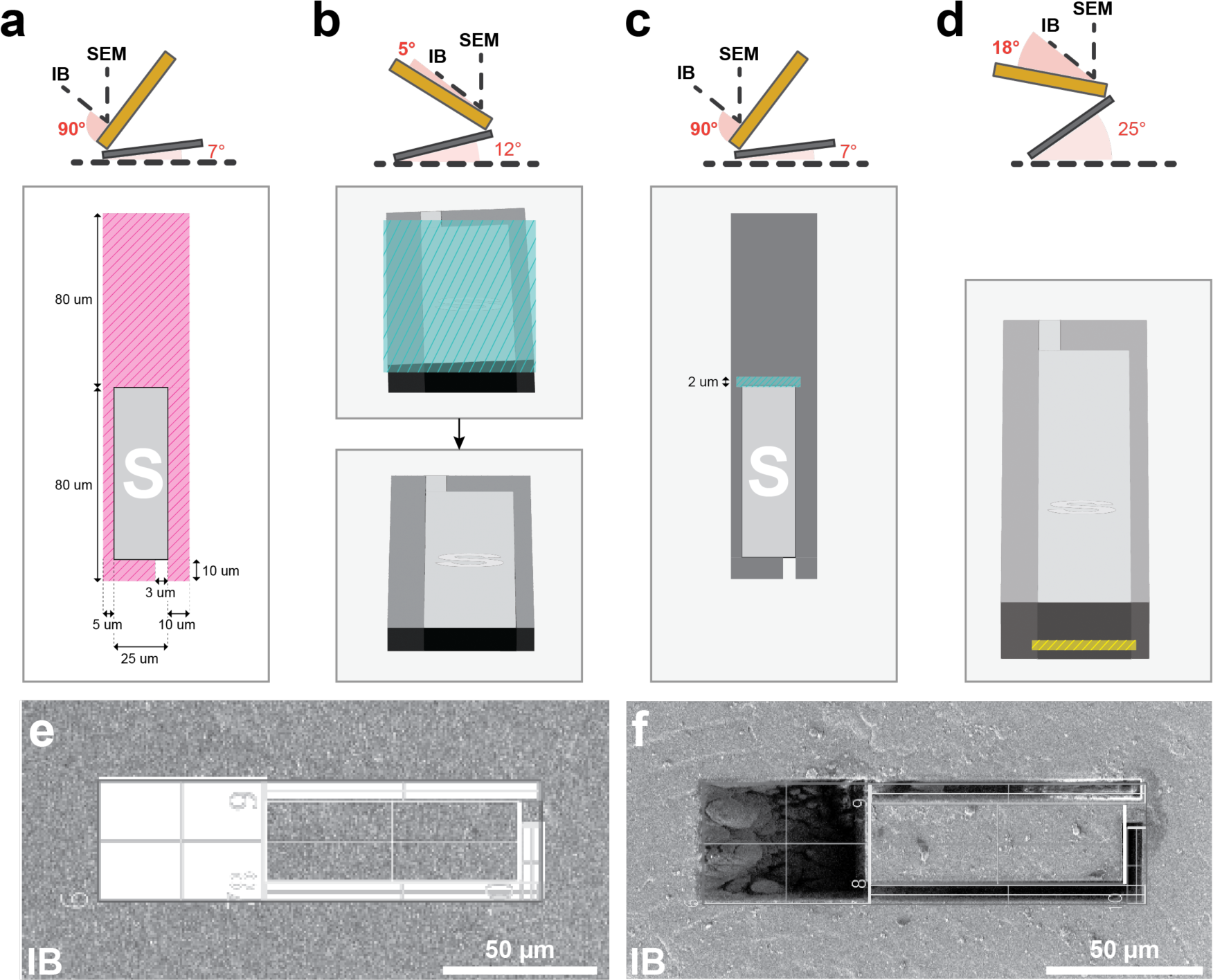
Sample site preparation for LO from the perpendicular direction. **a**, Sample trench preparation from the perpendicular position. **b**, Surface cleaning for needle attachment from the milling position at 12° stage tilt. **c**, Front edge polishing from the perpendicular position. **d**, Undercut milling from the milling position at 25° stage tilt. **e**,**f**, FIB image of the frozen sample with the patterns used to prepare the chunk site from perpendicular position before (**e**) and after milling (**f**). *Note. For illustration, images e and f are rotated 90° clockwise from the FIB view*.

**Supporting Figure 8.**
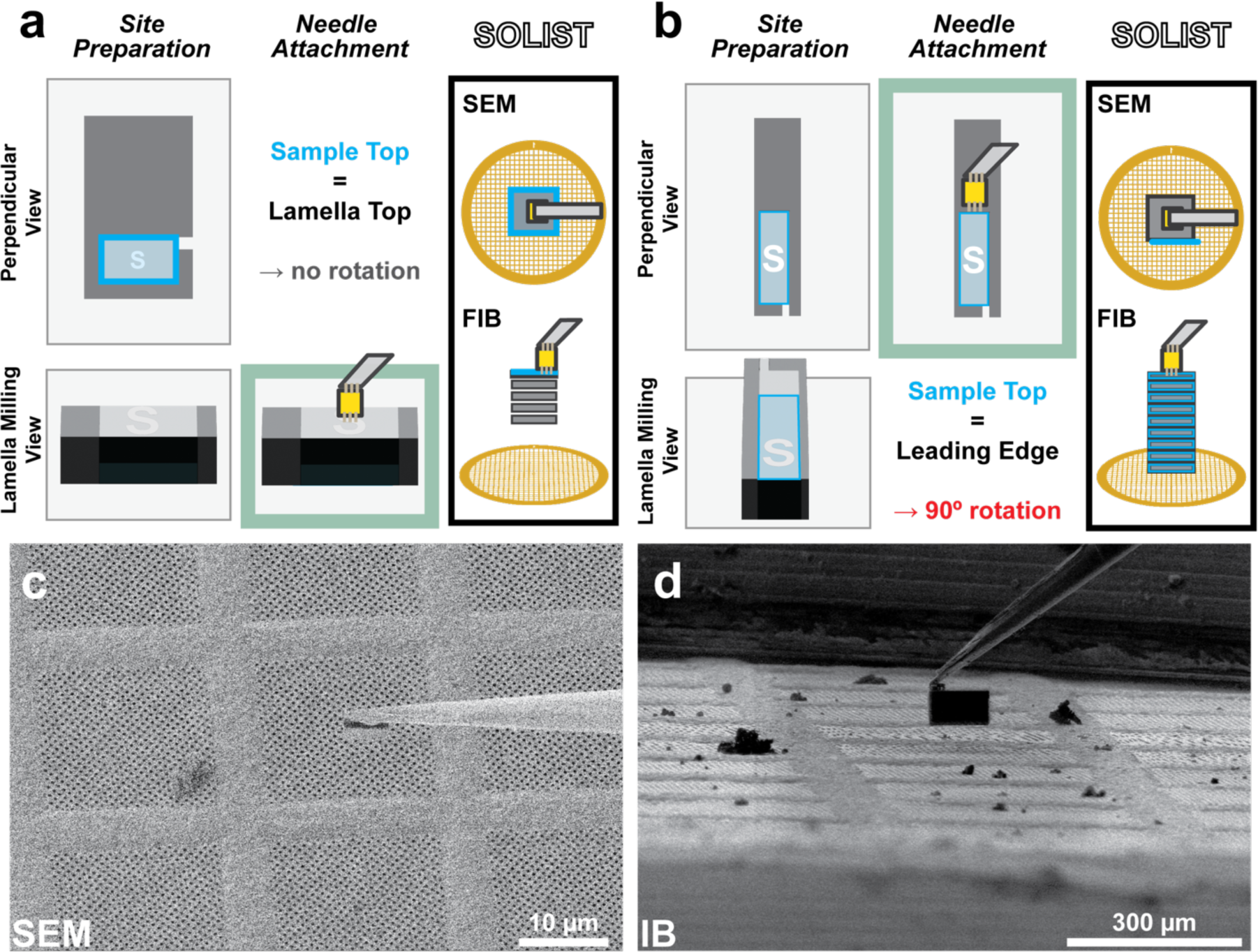
SOLIST lift-out procedure. **a,** For LO from the milling direction, the sample orientation remains unchanged, i.e., the chunk top (light blue) corresponds to the lamella top. To this end, the needle is inserted with the stage in milling position at 12° stage tilt and attached to the top of the sample chunk. **b,** For the LO procedure from the perpendicular direction, the sample is effectively re-oriented by 90°. Hence, the chunk top surface (light blue) becomes the lamella leading edge. Here, the needle is inserted with the stage close to the perpendicular LO position at 18° stage tilt and attached to the front edge. **c,d** SEM (**c**) and FIB (**d**) images of the Tungsten needle with the sample chunk attached to the gold adaptor and inserted over the LI grid.

## Online Methods

### SAMPLE PREPARATION

#### Organoids

Forebrain organoids were generated from H9 embryonic stem cells (ESCs) as previously described^1^. Embryoid bodies (EBs) were generated from ESCs colonies incubated in neural induction medium consisting of 2 μM Dorsomorphine (Stem cell technologies) and 2 μM A83-01 (Stem cell technologies). From day 7 to day 14, EBs were embedded in Matrigel (Corning) and patterned towards a forebrain fate in a medium consisting of DMEM:F12 (Gibco), 1X N2 Supplement (Life Technologies), 1X Penicillin/Streptomycin (Euroclone), 1X Non-essential Amino Acids (Gibco), 1X GlutaMax (Gibco), 1 μM CHIR99021 (Tocris), and 1 μM SB-43154 (Stem cell technologies). On day 14, embedded organoids were mechanically removed from Matrigel and incubated in differentiation medium composed of DMEM:F12, 1X N2 and B27 Supplements (Life Technologies), 1X Penicillin/Streptomycin, 1X 2-Mercaptoenthanol (Gibco), 1X Non-essential Amino Acids, 2.5 μg/ml Human Insulin (Sigma-Aldrich) until day 35. From day 35 to day 70, the differentiation medium was supplemented with 1% Matrigel. From day 70 onwards, organoids were incubated in maturation medium composed of Neurobasal medium (Life Technologies), 1X B27 Supplement, 1X Penicillin/Streptomycin, 1X 2-Mercaptoenthanol, 0.2 mM Ascorbic Acid (Sigma-Aldrich), 20 ng/ml BDNF (Peprotech), 20 ng/ml GDNF (Peprotech), and 1 μM Dibutyryl-cAMP (Stem cell technologies). Brain organoids were extracted from Matrigel as described by Qian X. *et al* (2020)^1^. They were then embedded in low-melting-point agarose (Invitrogen) and sectioned to 75 μm thick slices using a VT 1200S vibratome (Leica Microsystems).

#### *In vitro* chromatin sample

Condensation of chromatin droplets was performed *in vitro* as described previously^2^. In short, reconstituted chromatin was mixed with eGFP-labelled BRD4 containing only two bromodomains (BD1-BD2) in a phase separation buffer (25 mM Tris, pH 7.5, 150 mM KOAc, 1 mM Mg(OAc)_2_, 5% glycerol, 5 mM DTT, 0.1 mM EDTA, 0.1 mg/mL BSA). Just before freezing, 10% Ficoll 400 (Sigma-Aldrich) was added to the solution as cryo-protectant. To confirm droplet formation, the sample was observed on a Zeiss LSM980 confocal microscope with a 63x oil objective and using an excitation wavelength of 488 nm.

### HIGH-PRESSURE FREEZING

#### Planchette preparation

Before use, the planchettes were cleaned by sonication for 5 min, 30s on, 30s off, 40% Amplitude on a Branson SFX 550 (Thermo Fisher Scientific). Type B lids were polished with 1 μm lapping paper (Leica Microsystems) and then coated with 3 μl 0.1% soy lecithin (Sigma-Aldrich) in chloroform (Thermo Scientific).

#### Organoids

Just before freezing, slices were stained with Hoechst dye (Invitrogen) to later localize the cells embedded in the HPF buffer. They were then loaded in 100 μm-depth type A wells and frozen on a Leica EM ICE with 20% dextran (Sigma-Aldrich) and 5% sucrose (Sigma-Aldrich) supplied in the freezing medium.

#### *In vitro* droplets

The assembled condensation reaction with Ficoll 400 was frozen on a Leica EM ICE in 3 mm type A and B HPF planchettes (Leica Microsystems) by applying ∼0.8 μL of the solution to the 100 μm well of the type A before freezing.

### PLANCHETTE PLANING (OPTIONAL)

Copper HPF planchettes were cryo-planed using Leica EM UC7 ultramicrotome (Leica Microsystems) operated at −160°C. Approximately 30 µm of frozen buffer was removed. Rough trimming was performed with a TRIM45 diamond blade to remove the copper of the planchette (feed of 200 nm, speed of 60 mm/s). The obtained blockface was further trimmed with a separate TRIM45 blade for fine trimming with a feed of 50 nm and a speed of 30 mm/s.

### PLANCHETTE FLUORESCENCE MICROSCOPE PRE-SCREENING (OPTIONAL)

HPF planchettes were imaged at the cryo-fluorescence light microscope Thunder (Leica Microsystems) with a 50x / 0.9 NA objective and equipped with a cryo-stage. A customized adaptor was used to fit the HPF planchette into the Thunder loading cartridge. Planchettes overviews in reflected light, green (GFP) and blue (Hoechst) channels were acquired to define regions enriched with target structures.

### LIFT-OUT AND SOLIST PROCEDURE

Lift-out was performed at the identified areas on an Aquilos 2 cryo dual-beam FIB/SEM microscope equipped with the EasyLift lift-out system (Thermo Fisher Scientific)^3^. In general, milling position refers to the shuttle orientation for regular on-grid FIB milling and is defined as relative rotation 0°. The perpendicular position refers to a 180° relative rotation from the milling position.

#### Adaptor chunk preparation

An adaptor gold chunk of ca. 5-10 µm x 5 µm x 6 µm was prepared from a gold EM grid (UltrAuFoil, Quantifoil) following the procedure described previously and attached to the Tungsten needle of the EasyLift system^3^. Gold was chosen over copper for three reasons: 1) the chunk could directly be prepared from the loaded all gold support grid; 2) the milling of gold is easier and faster than copper; 3) sputter rates are higher for gold, and it is therefore easier to micro-weld the biological material to the LO needle. With the stage in the perpendicular position (stage tilt = 7°, relative stage rotation = 180°), 4 µm of material was removed from each side with regular cross-section patterns used at 5 nA and maintaining a 2 µm piece on the right side as attachment (Supporting Fig. 5a). The surface of the chunk was briefly cleaned from the milling position at an angle of 12° stage tilt using 300 pA with a cleaning cross-section pattern. An undercut was milled 5 µm below the surface with a regular cross-section pattern of 1 µm in *y* using 300 - 500 pA. The same current was used to clean 2 µm of material on the sides of the chunk, leaving the top 1 µm of the right side to connect the piece to the bulk (Supporting Fig. 5b). The stage was rotated back to the perpendicular position and a regular cross section pattern was used at 300 pA to clean material redeposition in the back (Supporting Fig. 5c). The needle was inserted above the EM grid with the stage in milling position at an angle of 12° stage tilt and lowered to enter in contact with the surface of the chunk. The needle was attached by milling micro-stitches on the gold and the remaining material on the left was milled with a regular cross-section pattern at 500 pA (Supporting Fig. 5d).

##### Sample site preparation

**1) For LO from the milling direction.** The Aquilos 2 stage was rotated so that the gallium ion beam was perpendicular to the surface of the sample (perpendicular position: stage tilt = 7°, stage rotation = 180°). A 40 µm-long area was cleaned in the front of the sample site and 5 µm-wide trenches were milled on the remaining sides of the chunk with 3 nA to 5 nA currents and regular cross-section patterns. A 3 µm piece was preserved on the left side to guarantee the attachment to the bulk (Supporting Fig. 6a).

The surface of the sample chunk could be cleaned from the milling position at an angle of 12° and with 300 pA using a cleaning cross-section pattern (Supporting Fig. 6b). The same current was used to clean the leading edge from the perpendicular position (Supporting Fig. 6c).

An undercut was milled 20-25 µm below the chunk surface from the milling direction at a high angle (∼ 35°) with 500 pA with regular cross-section pattern. At the same time, the sides of the sample chunk were cleaned, preserving 3 µm of sample on the top of the left side for connection (Supporting Fig. 6d).

**2) For LO from the perpendicular direction.** The sample site was prepared from the perpendicular position defining a sample trench of 25 µm in *x* and 80 µm in *y*. As for the LO from milling direction, the sides of the trench were cleaned with currents of 3 nA to 5 nA using regular cross-section patterns. Ca. 80 µm of material was cleaned in the front of the trench and 5 µm to 10 µm were removed from the other sides. A 3 µm piece was preserved on the rear side as attachment (Supporting Fig. 7a).

In case the surface was not homogeneously flat, the sample chunk was cleaned from the milling position at stage tilt of 12° and with 500 pA – 1 nA using a cleaning cross-section pattern (Supporting Fig. 7b). The same current was used to clean the front edge from the perpendicular position to improve the attachment of the needle (Supporting Fig. 7c).

An undercut was milled ∼10 µm below the chunk surface from the milling direction at 18-25° with a regular cross-section pattern at 1 nA. (Supporting Fig. 7d).

##### Lift Out

**1) LO from the milling direction.** The needle with the adaptor chunk was inserted with the stage in milling position at 12° stage tilt and lowered until it touched the sample surface without exerting significant force. Four micro-stitches of 0.8 µm x 1 µm were cut on the gold chunk with cleaning cross sections, milling direction towards the top, z dimension equal to 0.5 µm and currents set to 300 pA. The patterns were cut twice or three times, monitoring the gold redeposition between the sample and the adaptor chunk. When material started to redeposit, the patterns were backed up slightly from the sample and milled again.

To release the connection to the bulk sample, the remaining bridge was milled with a regular cross-section patterns at 500 pA. As soon as the sample chunk was free from the bulk, it was lifted out, rising the needle ∼50 µm above the planchette surface, before retracting the needle.

For a detailed illustration see Supporting Fig. 8a.

**2) LO from the perpendicular direction.** The needle with the adaptor chunk was inserted with the stage in perpendicular lifting-out position (stage tilt = 15°, relative stage rotation = 180°), lowered until it touched the sample front edge and attached to the trench as described above. The rear bridge connecting the sample to the bulk was milled with 500 pA using a regular cross section pattern, and the trench lifted out. For a detailed illustration see Supporting Fig. 8b.

##### SOLIST procedure

With the stage in the milling position, the needle with the sample chunk was inserted above a gold EM grid (UltrAuFoil, R2/2, 200 mesh, Quantifoil) used as support to receive individual lift-out sections and loaded in position 2 of the Aquilos2 shuttle. The needle was lowered until the sample chunk reached the grid foil and slightly backed up again to avoid stress on the gold film (as apparent from bending of the film).

For the correlative workflow, sections of 3-4 µm were cut but minimal sections of ca. 2 µm can be safely attached to the grids. A regular cross section pattern at 300-500 pA of at least 1 µm in *y* direction was placed on the sample chunk to release the section. This procedure was repeated until the LO chunk was exhausted.

With all the sample sections attached, the stage was rotated to the perpendicular position and several micro-stitches of 1.0 µm x 0.5 µm x 0.25 µm were placed on the left and right side of the sample and milled with cleaning cross-sections at 30 pA. The milling direction was set towards the center of the lamella.

The leading edge of each coarse lamella was cleaned from the same stage position with a cleaning cross-section pattern and z depth set to 1 µm at 300 pA.

The stage was rotate to the lamella milling position and the surface of each section could be cleaned from 12-15° using a cleaning cross-section pattern at 300 pA.

The grid was GIS-coated for 20 s from the lamella milling direction and 20 s from the perpendicular position before unloading (for storage or correlative workflow) or fine milling.

#### Fine milling

Lamellas were symmetrically thinned down to 800 nm with regular cross-section patterns at 300 pA and to 300 nm at 100 pA. Afterwards, rectangular patterns were used at 30 – 50 pA to reach 200 nm final pattern offset. To ensure homogeneous thickness along the lamella, the stage was tilted 1° up with respect to the used milling direction and a rectangular pattern at 30 - 50 pA was used to progressively mill from the rear towards the front of the lamella (overtilt milling).

#### Support grid preparation for the correlative workflow

The support grid (UltrAuFoil, R2/2, 200 mesh, Quantifoil) was plasma-cleaned at 30 mA for 30 s (GloQube Plus Glow discharge system, Quorum Technologies). Autofluorescent beads (Dynabeads MyOne, Life Technologies-Thermo Fischer Scientific) were washed following manufacturer instructions and diluted in deionized water to obtain a proper distribution on the grid. From this suspension, 3 µL of beads were applied to the grids, and after buffer evaporation the grid was clipped in a FIB-type autogrid and loaded into the microscope shuttle (Position 2).

#### Cryo-fluorescence light microscopy

Fluorescent image stacks of the coarse lamellas were acquired on a cryo-confocal microscope (Stellaris 5, Leica Microsystems) equipped with 50x / 09 N.A. objective and a cryo-stage. The pixel size was 216 nm (*x,y*) and the *z*-step was 330 nm. Images were acquired using an excitation wavelength of 630 nm and 489 nm for reflected light and eGFP-labelled protein, respectively. The bead signal was acquired separately in the same channel of eGFP using a higher intensity level.

#### Correlative fine milling

Support grids were transferred to the Aquilos 2. A FIB image of the sample section was acquired at 12° stage tilt and in the milling position and was registered to the fluorescence data with the 3D Correlative Toolbox (3DCT) software as described previously^4, 5^. In brief, “gaussian fit” was used to identify the x, y, and z position of five to ten beads in the eGFP channel acquired at high intensity. The same beads were identified in the FIB view to calculate a transformation matrix. Then, the target structures were identified in the eGFP channel which had been recorded at lower intensity. Their z position was fit by the software (“Gaussian Fit z”) and predicted on the FIB view. Lamellas of ca. 200 nm (final pattern distance) were cut at the predicted sites. Regular cross-sections were used at 300 pA to reach a thickness of 800 nm and then at 100 pA until 300 nm. Rectangular patterns at 50 pA or 30 pA were used to reach the final thickness of ∼ 180-200 nm (final pattern offset).

### CRYO-ELECTRON TOMOGRAPHY

#### Data acquisition

The SOLIST grids with the sample sections were loaded on a Titan Krios G4i operated at 300 kV equipped with Falcon4i direct electron detector and SelectrisX energy filter (ThermoFisher Scientific). For the correlative workflow, 3DCT was used to register fluorescence data to SEM view of the final lamella or to TEM montage to guide correlative TEM acquisition^4, 5^. Cryo-tomograms were recorded at 1.5 Å/pix (81k x magnification) for the *in vitro* LLPS droplets and at 1.962 Å/pix (64k x magnification) for the brain organoids. The defocus range was set to −1 μm to −3 μm. A dose-symmetric tilting scheme with an angle increment of 3° was used from −60° to +60° with respect to the lamella pre-tilt. Tilt series were acquired using the SerialEM tilt series controller or PACEtomo and 3 e/Å^2^/tilt (total dose for 41 images: 123 e/Å^2^)^6, 7^.

#### Data processing

Tilt series were processed in TOMOgram MANager (TOMOMAN) [https://github.com/williamnwan/TOMOMAN.git]. Frame alignment of the EER data was performed in Relion4 with EER groups adjusted to 0.5 e/Å^2^ ^8^. The final tomograms were denoised using cryoCARE and deconvolution-filtered to enhance the signal of biological-relevant structures^9^.

Automated membrane segmentations were performed with tomosegmenTV^10^. For animations, these segmentations were manually adjusted in AMIRA (Thermo Fisher Scientific) and animated in ChimeraX^11^.

#### Figure Composition

Figures were created in Adobe Illustrator and in Figure 2 and Supporting Figure 3 assets of biorender.com were used.

